# Evolution in novel environments: do restored prairie populations experience strong selection?

**DOI:** 10.1101/824409

**Authors:** Susan M. Magnoli, Jennifer A. Lau

## Abstract

When populations colonize new habitats, they are likely to experience novel environmental conditions, and as a consequence may experience strong selection. While selection and the resulting evolutionary responses may have important implications for establishment success in colonizing populations, few studies have estimated selection in such scenarios. Here we examined evidence of selection in recently established plant populations in two prairie restorations in close proximity (< 15 km apart) using two approaches: 1) we tested for evidence of past selection on a suite of traits in two *Chamaecrista fasciculata* populations by comparing the restored populations to each other and their shared source population in common gardens to quantify evolutionary responses and 2) we measured selection in the field. We found evidence of past selection on flowering time, specific leaf area, and root nodule production in one of the populations, but detected contemporary selection on only one trait (plant height). This demonstrates that while selection can occur in colonizing populations, resulting in significant evolutionary responses in less than 6 generations, rapid evolutionary responses may be weak in even nearby populations sown with the same source population. Because contemporary measures of selection rarely predicted observed evolutionary responses, it also suggests that selection likely differs over the early stages of succession that characterize young prairies.

## Introduction

We often assume that colonizing populations will experience strong selection if they encounter novel biotic and abiotic conditions in their new habitats. But while strong selection may drive the rapid phenotypic changes observed in some colonizing populations (reviewed in Bossdorf et al. 2005; Colautti et al. 2009; Felker-Quinn et al. 2013), few studies have actually estimated selection in such scenarios (Colautti and Lau 2015), calling into question the validity of this assumption. Furthermore, studies of selection on invasive vs. native populations and anthropogenically disturbed vs. undisturbed populations (where we also expect to find strong selection) show mixed results. For example, a synthesis of studies comparing selection gradients in invasive vs. native populations detected no evidence of stronger selection gradients in invasive populations (although selection differentials which include indirect selection acting on correlated traits were significantly stronger for invasive populations) (Colautti and Lau 2015). Similarly, a recent meta-analysis examining the strength of selection on populations experiencing anthropogenic disturbance, which included a small number of introduced populations, Fugère and Hendry (2018) found that selection was generally not stronger in human-disturbed environments.

The lack of evidence in support of the hypothesis that populations in novel and disturbed environments are under strong selection is surprising, if we assume that populations are optimally adapted to their native environments and that environmental change or introduction to a new environment pushes them away from that adaptive peak. However, there are at least two potential broad explanations for these patterns. First, selection actually might not be particularly strong in novel environments if the novel environment increases mean fitness, if the populations colonizing that environment are pre-adapted to novel conditions, or if there is no genetic variation in responsiveness to the new environmental conditions (i.e., all genotypes within a population experience increased or decreased fitness in the novel environment to the same extent). For example, strong selection will not occur if populations face decreased opportunity for selection when environmental conditions in a new habitat increase the mean or decrease the variance of absolute fitness (Crow 1958; Caruso et al. 2017). This could occur if a population colonizes a habitat with greater resource availability than its home site, and would weaken the strength of selection even in the presence of novel environmental conditions (Fugere and Hendry 2018). Strong selection also may not occur if successful colonizers are pre-adapted to environmental conditions in their new habitats (that is, environmental conditions in the new habitat are similar to those in the native habitat). Alternatively, even if the environment is quite different, strong selection will not result if the same traits promote fitness in both the novel and historical environments. Pre-adaptation is often hypothesized to be a factor in successful species invasions (Cadotte et al. 2018), and if it is common this could help explain a lack of strong selection in some invasive populations. Finally, in some cases, selection may not be particularly strong in a novel environment if there is little genetic variation in fitness or trait responses to that environment. For example, although novel environments are often hypothesized to lead to the expression of cryptic genetic variation (Gibson and Dworkin 2004; Paaby and Rockman 2014), which should increase the likelihood of detecting strong selection, in an *Arabidopsis thaliana* recombinant inbred line population, no genetic variation in CO_2_ responsiveness was detected. As a result, dramatically elevated atmospheric CO_2_ concentrations that the populations had never historically experienced failed to alter natural selection on any traits measured (Lau et al. 2007).

Second, colonizing populations may experience strong selection, but we may miss it if it occurs rapidly. This may be especially likely during colonization events, as selection on some traits may be strong during establishment, but weaken or become more similar to patterns of selection in the native range over time. For example, in invasive *Alliaria petiolata* populations in North America, production of high amounts of phytotoxins may be selectively favored under initial invasion conditions with high interspecific competition, but populations evolve lower phytotoxin production over time as intraspecific competition increases (Lankau et al. 2009). If scenarios such as this commonly occur, whether or not strong selection is detected in colonizing populations may depend on when it is measured.

Ecological restorations present a unique opportunity to examine selection on colonizing populations. In many restorations, plant populations are seeded into highly-anthropogenically disturbed environments, along with other populations with which they may not share a recent evolutionary history. Populations are unlikely to be optimally adapted to these novel abiotic and biotic conditions, which could generate strong selection for evolutionary change (Rice and Emery 2003; LaRue et al. 2017). In addition, unlike most invasive species, we know when populations were planted in restorations, and can even save seed from the original source populations, which allows us to examine selection in two ways. First, we can compare restored populations to their original source populations in common gardens to look for evidence of rapid evolutionary changes in the time since restoration, which would suggest strong selection had occurred in the recent past. Second, we can estimate selection directly in the field with phenotypic or genotypic selection analyses (Lande and Arnold 1983; Rausher 1992). In this study we use both of these approaches to estimate selection on a suite of traits in annual plant populations in two recently restored prairies.

## Methods

### Study System

*Chamaecrista fasciculata* Michx. (hereafter *Chamaecrista*) is an annual legume native to eastern North America found in prairies and other disturbed sites. *Chamaecrista* is commonly planted in prairie restorations (Grman et al. 2015), where it forms facultative mutualistic interactions with nitrogen-fixing rhizobium bacteria, which provide plants nitrogen in exchange for carbon, provided compatible rhizobium strains are present. In 2010, two former agricultural fields (one 13 hectare and one 11 hectare) near Kellogg Biological Station in southwest Michigan, Lux Arbor (42°28’23” N, 85°26’50” W) and Marshall (42°26’37” N, 85°18’34” W), were planted with identical prairie seed mixes (containing 19 grass and forb species, including *Chamaecrista*, seeded at 0.28 kg/ha, roughly 26,700 seeds/ha) as part of a large bioenergy experiment being conducted by the Great Lakes Bioenergy Research Center (https://www.glbrc.org/). Biomass from each prairie is harvested every year using identical protocols (see Stahlheber et al. 2016 for full site details). Seeds were saved from the original seed mix.

We predicted that the original source *Chamaecrista* population may not have been optimally adapted to restoration sites in southwest MI; therefore, these populations might experience strong selection on phenological and morphological traits during population establishment, resulting in genetic differences between extant populations and the original source seeds. Our knowledge of the growing conditions of the original source population is somewhat limited, but we know seeds were produced on a seed farm in Houston County, MN (Shooting Star Native Seeds, *pers. comm.*), which is 1-2° higher latitude than our restoration sites in southwest MI, with slightly lower average rainfall. This latitude difference could lead to a mismatch in optimal phenology between the source and restoration sites, as higher-latitude populations of *Chamaecrista* have been shown to flower earlier than lower latitude populations (Etterson and Shaw 2001). In addition to the abiotic differences between the source and restoration sites, there are likely biotic differences that could lead to evolutionary changes in restored populations. Seed harvesting and propagation for restoration can lead to evolutionary changes in source populations (McKay et al. 2005; Vander Mijnsbrugge et al. 2010; Dyer et al. 2016; Gallagher and Wagenius 2016; Ensslin et al. 2018) that may produce populations that are poorly adapted to diverse communities. We therefore might expect traits related to competitive ability, such as plant height, to be under selection in diverse restoration sites. Similarly, light availability for a relatively short-statured plant like *Chamaecrista* is likely to differ between a monoculture and a diverse prairie community, which could lead to selection on traits like specific leaf area. Microbial communities are also likely to differ between the source and restoration sites (Koziol et al. 2018), which could influence traits related to *Chamaecrista’s* mutualism with nitrogen-fixing rhizobia.

### Evidence of past selection and estimates of contemporary selection

To determine whether these recently established *Chamaecrista* populations have undergone evolutionary change, and to estimate selection, we conducted a field experiment using seeds collected from the two extant populations, as well as resurrected seeds saved from the original seed mix. In September 2015 (likely six *Chamaecrista* generations after the populations were planted, as *Chamaecrista* has a limited seed bank (Fenster 1991)) we collected 5-20 seeds from each of 100 individuals from each site. To do this we established five 100 meter transects at each site, and collected seeds from the nearest *Chamaecrista* individuals to the transect at five-meter intervals. We grew seeds from these plants, along with seeds from the original seed mix (seeds had been stored mixed with all other species in the original seed mix in plastic mesh bags at room temperature since 2010), in the greenhouse for one generation to minimize maternal effects. For the two extant populations, we grew one seed from each of 96 of the 100 maternal plants. Each plant was randomly assigned to be a sire or a dam, and pollen from each sire was used to pollinate two dams, for a total of 64 full-sibling families (32 half-sibling families) per site. We did not include family structure when pollinating flowers of the original source plants, due to low germination of the original source seed (7%; in contrast, approximately 95% of seeds collected from the two extant population germinated). Instead we used one plant as a pollen donor on a given day (for approximately 60 days), so that every plant was crossed with every other plant several times.

In May 2016 we germinated seeds produced by these plants in the greenhouse (we had approximately 95% germination success and no differences in germination among the three populations). We transplanted seedlings into randomized locations within six 4 × 4m plots (each divided into 16 1 × 1m subplots with seedlings spaced 16cm apart) in each prairie site {(2 seedlings/extant population full-sib family × 64 families/extant population × 2 extant populations + 64 original source population seedlings) × 6 plots × 2 sites; N=3840 total seedlings}, in a reciprocal transplant design. We grew all populations at both sites because if populations had diverged in trait values, including all populations would expand the phenotypic distribution and allow for a better estimate of the overall fitness function in our selection analyses (Conner and Hartl 2004). We fenced half of the plots to exclude deer and small mammals, to ensure that herbivores did not kill all of the seedlings (in a previous experiment, small mammal herbivory driven by a vole outbreak led to 95% plant mortality); however we detected little herbivore-induced mortality in this experiment. We disturbed existing vegetation as little as possible when planting seedlings.

Over the course of the growing season we monitored survival and recorded day of first flower. In July 2016 we collected the third fully-expanded leaf from the top of each plant to measure specific leaf area (SLA). In September 2016, when most fruits were mature enough to count seeds, we harvested all aboveground biomass, measured plant height, and counted the number of seeds produced by each plant as a lifetime female fitness measure. We also harvested a root sample from each plant by digging up the top 5-10cm portion of the root, and then counted the number of root nodules to estimate the number of root nodules/length of root.

### Past selection - evidence of evolutionary change

To determine whether the two extant populations differ from each other or their original source population, we compared trait values from each population in the 2016 common garden experiment using linear mixed models in the lme4 package (Bates et al. 2015) in R v. 3.5.1 (R Core Team 2018). Trait values (date of first flower, nodules/cm root, SLA, and height) were included as response variables. Source population (Lux, Marshall, Original), site (Lux, Marshall), fencing (present, absent), and all interactions between these variables were included as fixed effects, and plot nested within fencing, and subplot nested within plot were included as random effects. For the height model, we excluded plants that had been browsed by herbivores. We log transformed response variables when necessary to better fit model assumptions. Because the root nodule data was both zero-inflated and right-skewed, we first used a logistic regression model with whether a plant nodulated (yes/no) as the response variable. Then we used a linear model with the number of nodules/cm root as the response variable, including only the plants that formed nodules. We tested significance using type III sums of squares in the ANOVA function in the *car* package (Fox and Weisberg 2011) with sum contrasts, and calculated least squares means and conducted Tukey’s post-hoc multiple comparisons tests using the *emmeans* package (Lenth 2018).

Due to low germination of the original source seed (7%), there was a risk that the mean trait values we found for this population were not accurate representations of the entire source population. A biased sample could result if the measured traits are correlated with seed viability (Franks et al. 2019), or if the small sample size led to founder effects. To estimate whether founder effects influenced our conclusions, we bootstrapped trait distributions for each trait in the extant populations by repeatedly drawing seven families at random from each population, to calculate a distribution of population mean trait values controlling for sample size. When we found trait differences between either extant population and the original source population in the experiment, we compared where the original source mean fell on the bootstrapped extant population trait distributions. In all cases, the original source mean fell outside the 95% confidence intervals of the distributions, indicating that trait differences were likely not the result of a founder effect in our sample of the source population (Fig. S1). However, we cannot rule out biases caused by trait correlations with long-term seed viability. Therefore, while we present traits means for the original source population in the results, we focus on differences between the two restored populations as strong evidence of past evolutionary changes.

### Contemporary selection - genotypic selection analyses

To estimate the strength and direction of selection we conducted selection analyses on *Chamaecrista* at both sites using linear models. These analyses regress relative fitness on standardized trait values to estimate the strength and direction of selection on measured traits (Lande and Arnold 1983). We used family mean trait and fitness values to conduct genotypic selection analyses, which remove biases caused by environmentally-induced covariances between traits and fitness (Rausher 1992). We used data from the two extant populations only (because we had no family structure for original source plants), and included only plants from unfenced plots as we wanted estimates of selection on traits under natural field conditions. We excluded height measurements from plants that had been browsed by deer from the analysis because we could not obtain accurate measurements for these individuals. These analyses included all plants grown at each site, regardless of population (that is, we combined data from Lux plants grown at the Lux site with Marshall plants grown at the Lux site to examine selection at the Lux site).

We tested whether selection differed across sites, by conducting ANCOVA with the residuals of family mean relative fitness as the response variable and standardized trait values (height, flowering time, root nodules, SLA) and their interactions with site as predictor variables. We used the residuals of relative fitness after the effects of plot had been removed to reduce the influence of spatial variation. To estimate the strength and direction of selection on traits at each site we ran separate models for each site, similar to the one described above but without the trait x site interaction terms. To examine non-linear selection coefficients and correlational selection, we used models with traits, quadratic terms (traits squared), and all trait cross-products as predictor variables (quadratic coefficients were doubled (Stinchcombe et al. 2008)).

### Contemporary selection - phenotypic selection analyses

To explore selection on traits in different years, we also conducted phenotypic selection analyses on *Chamaecrista* trait and fitness data collected in 2014 and 2015 (we used phenotypic and not genotypic selection because we had no family structure for individuals measured in these years). We censused 100 and 200 plants from each site in 2014 and 2015, respectively (sampled along transects similar to those described above for seeds collections), and measured plant height, specific leaf area (2014 only), date of first flower (2015 only), and counted seeds to estimate fitness. Because different traits were measured in each year, we calculated linear selection gradients using separate models for each year. As with the genotypic selection analysis, we regressed trait × site interactions on relative fitness to determine whether selection differed between sites, then ran separate models for each site without the site interactions to estimate direct selection. As described above we included quadratic terms, and cross-products, to examine non-linear and correlational selection. To determine whether selection on height differed between years (the only trait we measured in both years), we conducted ANCOVA with relative fitness as the response, and height, year, and their interactions as predictor variables. The two sites were analyzed separately. These analyses test for differences in the height selection differential (as opposed to the selection gradient) across years. Selection differentials estimate total, rather than direct, selection acting on a trait.

## Results

### Past selection - evidence of evolutionary change

We found that populations differed significantly in flowering time and SLA. The Lux population flowered significantly earlier than the Marshall population, which flowered significantly earlier than original source plants, although this effect was only statistically significant when plants were grown at the Lux site (population × site: F_2,2074_=10.62, p<0.0001; Fig. 1). The Lux population also had significantly lower SLA and tended to be taller than original source plants (SLA: F_2,2095_=5.26, p=0.005; Height: F_2,2031_=2.74, p=0.06 Fig. 2a,b). We also found differences in nodule formation between populations. Lux plants were more likely to nodulate than Marshall plants regardless of site, but neither population differed significantly from the original source (population: χ^2^=8.43, p=0.01; Fig. 2c). However, of those that did produce nodules, Lux plants tended to produce more nodules than the original source population (17%) (population: F_2,1419_=2.84, p=0.058; Fig. 2d). All traits differed significantly between sites, suggesting that these traits are highly plastic, although with the exception of flowering time (see above) populations did not differ in plasticity (few significant source × site interactions; Table S1). Plasticity in height (the only trait under selection in 2016) appeared to be adaptive as plants were taller at Lux and selection favored taller plants at Lux, but not at Marshall.

**Figure 1.**
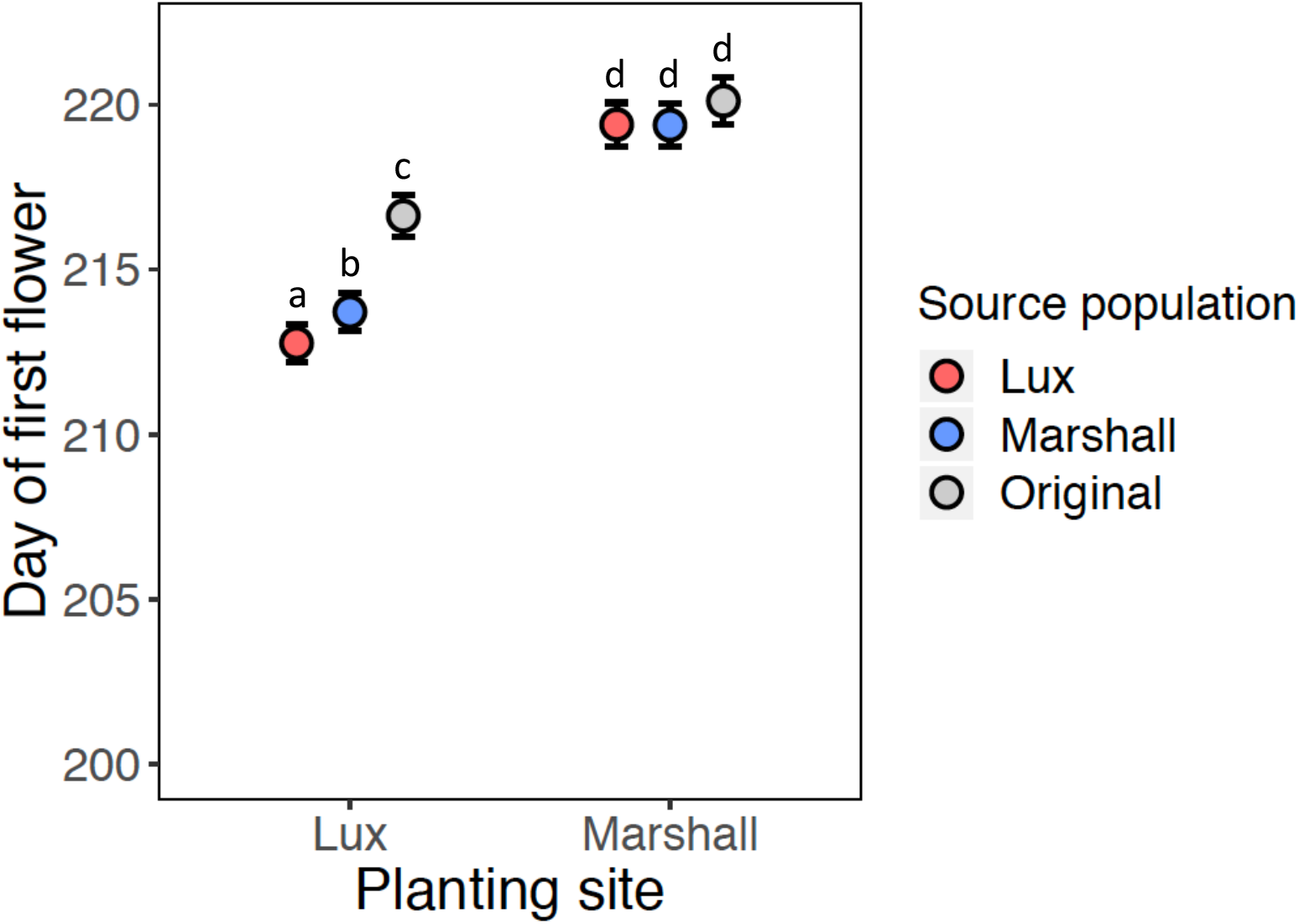
Population mean (estimated marginal mean±SE) day of first flower at both restoration sites.

**Figure 2.**
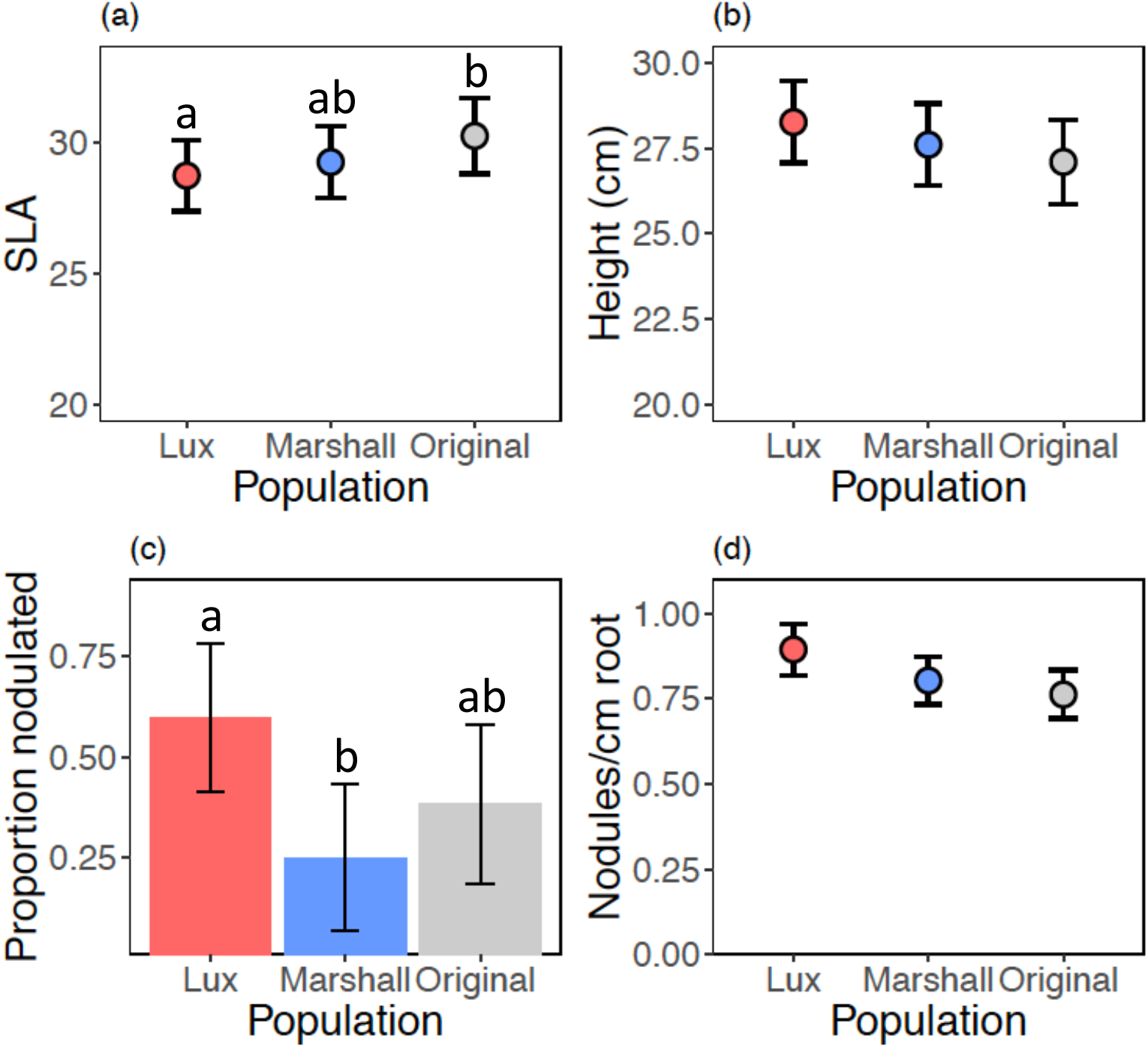
Population trait means (estimated marginal mean±SE) showing (a) specific leaf area (SLA), (b) height, (c) proportion of plants that formed root nodules, and (d) root nodules/cm root produced by plants that nodulated.

### Contemporary selection - genotypic selection analyses

Plants grown at the Marshall site had very low fitness compared to those grown at the Lux site, with many families not producing any seeds (Fig. S2). Selection overall did not differ between sites (site × trait interactions, all P > 0.18; Table S2). However, when we calculated selection gradients for each site, we found selection for taller plants at Lux (F_1,121_=7.42, p=0.007; Table 1) and no significant selection on height at Marshall (F_1,75_=0.09, p=0.77; Table 1). Although we detected some evidence for non-linear (stabilizing) selection on height at Lux (F_1,111_=17.90, p<0.001), the quadratic selection gradient was much smaller in magnitude than the directional selection coefficient (Table 1) and selection appeared linear although slightly weaker as height increased across most of the range of trait values (Figure S3). We found no significant directional selection on flowering time, root nodules, or SLA at either site (Table 1). There was no significant correlational selection on pairs of traits at either site.

**Table 1.** Directional and quadratic genotypic selection gradient (β) estimates ± SE at the Lux and Marshall sites in 2016. Quadratic estimates are in parentheses. \**P*<0.05

| | Lux $\beta$ | Marshall $\beta$ |
| --- | --- | --- |
| **Date of first flower** | -0.002±0.07 (0.12±0.12) | -0.05±0.27 (-0.04±0.52) |
| **Nodules/cm root** | -0.04±0.07 (0.12±0.12) | -0.16±0.26 (-0.02±0.48) |
| **SLA** | 0.04±0.08 (-0.12±0.10) | 0.07±0.29 (0.02±0.41) |
| **Height** | 0.25±0.07* (-0.16±0.06) | -0.09±0.31 (-0.52±0.64) |

### Contemporary selection - phenotypic selection analyses

We detected selection for increased height at both sites in 2014 (Table 2), although it was stronger at Lux (height × site F_1,179_=4.21, p=0.04), consistent with results from the 2016 genotypic selection analysis at Lux. We also detected evidence for significant non-linear selection on height (Table 2; Fig. S4a), such that the strength of selection increases as height increases. There was no significant selection on SLA at either site (Lux F_1,89_=0.03, p=0.86; Marshall F_1,90_=1.90, p=0.17). We detected a significant correlational selection gradient on height and SLA at Marshall, suggesting selection for higher SLA when plants are tall but selection for lower SLA when plants are short (F_1,87_=4.43, p=0.04; Fig. S5a).

**Table 2.** Directional, quadratic, and correlational phenotypic selection gradient (β) estimates ± SE at the Lux and Marshall sites in 2014 and 2015. Quadratic estimates are in parentheses. Note that selection was not measured on all traits in all years. \**P*<0.05, ** *P*<0.01, \*\*\**P*<0.0001

|  | 2014 |  | 2015 |  |
| --- | --- | --- | --- | --- |
| | Lux $\beta$ | Marshall $\beta$ | Lux $\beta$ | Marshall $\beta$ |
| **Height** | 0.73±0.10***<br>(0.36±0.08***) | 0.47±0.08***<br>(0.28±0.04***) | 0.42±0.08***<br>(0.20±0.06) | 0.92±0.12***<br>(0.72±0.16)*** |
| **SLA** | -0.02±0.09<br>(-0.14±0.10) | -0.10±0.07<br>(0.06±0.05) | -- | -- |
| **Height\*SLA** | -0.14±0.15 | 0.25±0.12* | -- | -- |
| **Date of first flower** | -- | -- | -0.03±0.08<br>(-0.08±0.04) | 0.05±0.13<br>(-0.02±0.18) |
| **Height\*Date of first flower** | -- | -- | -0.02±0.08 | 0.34±0.17* |

In 2015 we again detected selection for increased height at both sites (Table 2), but it was stronger at Marshall than Lux (height × site F_1,264_=20.98, p<0.001). We again found evidence that selection on height was non-linear, but quadratic coefficients were much smaller in magnitude than the directional selection coefficients and selection appeared to be primarily directional across the range of trait values included in this experiment (Fig. S4b). There was no significant selection on flowering time at either site (Lux F_1,145_=0.11, p=0.75; Marshall F_1,118_=0.12, p=0.73); however we detected some evidence for correlational selection gradients, suggesting that selection on flowering time depends on plant height. Specifically, at Marshall selection appeared to favor later flowering when plants were tall (flowering time × height: F_1,115_=3.85, p=0.052; Fig. S5b), although we detected no significant interaction between flowering time and height at Lux (F_1,142_=0.06, p=0.81). Comparing the strength of selection on height across years at each site, we found opposing patterns (height × site × year: F_1,449_=19.26, p<0.001). Although selection consistently favored taller plants, Lux selection on height was significantly stronger in 2014 (height × year F_1,237_=7.68, p=0.006) while at Marshall selection was stronger in 2015 (height × year F_1,212_=11.09, p=0.001).

## Discussion

Despite the expectation that colonizing populations are likely to experience strong selection as they encounter novel environmental conditions in new habitats, our results suggest that strong selection, when it occurs, may be transient. We found that two restored *Chamaecrista* populations differed from each other, and potentially from their original source population, in some traits. This suggests that at least one of these populations may have experienced selection strong enough to result in evolutionary changes in the six years since establishment. In contrast, we found little evidence of contemporary selection acting on these same traits in the year of our experiment. These results demonstrate that selection pressures on colonizing populations may vary through time and differ between identically-restored sites in close proximity.

### Past selection - evolutionary differences among populations

By comparing traits between the two restored populations and their original source population we detected evolutionary changes that indicate possible past selection. The Lux population flowered earlier than the Marshall and original source populations (when grown at the Lux site), was more likely to nodulate than the Marshall population and tended to make more nodules than the original source population, and had lower specific leaf area than the original source population. We need to be cautious when drawing conclusions based on comparisons to the original source population given low germination rates of the resurrected seeds, as seed longevity could be correlated with other traits, potentially biasing our results (Franks et al. 2019). However, the fact that the Lux population also differed from the Marshall population suggests that these populations have diverged over the past 6 years. Because the Marshall population did not differ significantly from the original source population in most traits, the most parsimonious explanation is that evolutionary changes have occurred in the Lux population but were minimal in the Marshall population. These changes are likely the result of selection and not drift, given that the Lux population is fairly large (>10,000 individuals, *pers.obs.*).

### Contemporary selection

While at least one of these restored populations may have experienced selection in the time between initial restoration and our study, we find little evidence of strong contemporary selection. Categorizing selection as strong or weak is dependent on the definition of these terms, which are relative (Conner 2001). However, we can compare our selection gradients to those in other studies to determine their relative strength. In a review of studies that estimated selection in natural populations by Kingsolver et al. (2001), the median |β| value for linear selection was 0.16. For three of our four measured traits we found no significant selection (that is, our gradients were not statistically significant from 0) and all estimates were relatively small in magnitude (< 0.16). In contrast, we found selection for increased height at both restoration sites (with the exception of the Marshall site in 2016) across three years, with β ranging from 0.36 to 0.92 (Tables 1, 2). Taken together, these results suggest that while selection on plant height is relatively strong in these populations, selection in general is not particularly strong.

### Temporal variation in selection and differences between sites

The mismatch we observe between past and current selection could be explained by several factors. First, it may be that we find evidence of past selection, but no contemporary selection, on some traits because they have already rapidly evolved to their evolutionary peak. In such a case, we might expect to observe stabilizing selection, but this was not the case for any trait in which we observed evolutionary changes (Table 1), although stabilizing selection can be difficult to detect (Conner 2001; Kingsolver et al. 2001). Second, and perhaps more likely, our findings may reflect variation in selection through time. Many long-term studies of selection have found temporal variation in selection, due to fluctuations in both abiotic and biotic environmental conditions (Siepielski et al. 2009). While yearly fluctuations in climatic conditions could lead to variation in selection, we might also expect selection on these traits to change over the course of restoration establishment/succession. For example, selection on *Danthonia spicata* traits in habitats at different successional stages differed in both the magnitude and direction of selection, depending on successional stage (Scheiner 1989). Prairie restorations tend to be dominated by weedy annuals in earlier years, followed by the establishment of perennial grasses and forbs in later years (Schramm 1990). These changes in plant composition during succession could lead to changes in competition and the abiotic environment that affect selective pressures on *Chamaecrista*, which may be likely given that *Chamaecrista* is an early-successional annual that often grows in recently disturbed habitats (Galloway and Fenster 2000).

### Conclusions

In this study, we demonstrate that plant populations recently established in anthropogenically-disturbed restoration sites can experience selection, but that it may vary over time and relatively small geographic distances. We show that two populations, originating from a shared source population and planted into identically-prepared restorations only 15km apart, can experience different selective pressures and/or differential responses to selection. While we find evidence of selection and/or evolutionary responses on all traits measured in the Lux population, there is less evidence of selection on the Marshall population. These differences could be a direct result of environmental differences between the two sites: despite their close proximity, plant community composition and some abiotic conditions (such as soil nitrogen and water holding capacity) differ (Stahlheber et al. 2016), which could generate different selection pressures at each site. Alternatively, we may find evidence of past selection in one population but not the other not because of differences in selection pressures, but because of differences in the populations’ capacities to respond to selection or the degree to which the populations were preadapted to local site conditions. Preadaptation is an unlikely explanation for why we find less evidence of selection in the Marshall population, given the very low fitness of plants at that site (Fig. S2). However, the two populations may differ in their capacities to adapt due to differences in population size; the Marshall population has been consistently smaller than the Lux population since initial establishment (Magnoli 2018). Smaller population size may lead to lower genetic diversity (Leimu et al. 2006), which may mean that the Marshall population is less able to respond to selection than the larger, more genetically diverse Lux population. Further exploration of the drivers and limits of adaptation in restored populations will give insight into the evolutionary processes underlying colonization success in degraded habitats.

## Acknowledgements

We thank S. Davis, M. Hammond, A. Hammond, T. Koffel, B. Motsa, N. Turley, J. Winters, and M. Zettlemoyer for help with field work, and L. Brudvig, J. Conner, M. Weber, and members of the Lau lab for providing comments that improved this manuscript. Support for this research was also provided by the US DOE Office of Science (DE-FCO2-07ER64494) and Office of Energy Efficiency and Renewable Energy (DE-ACO5-76RL01830) to the DOE Great Lakes Bioenergy Research Center, the NSF Long-term Ecological Research Program (DEB 1637653) at the Kellogg Biological Station, and Michigan State University AgBioResearch. This work was funded by a G.H. Lauff Summer Graduate Research Award awarded to S.M.M.

## Supplementary Material

**Table S1.**
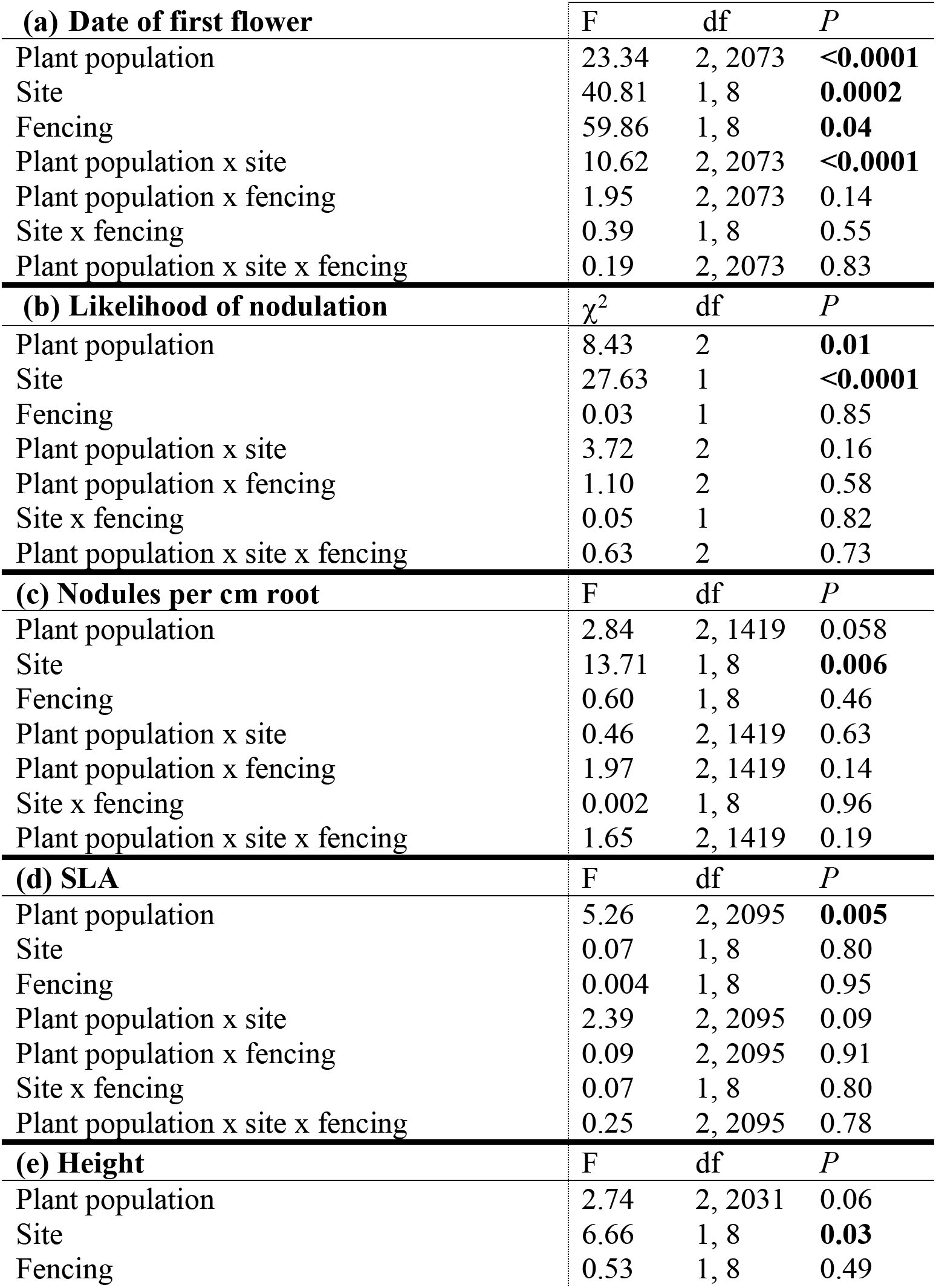
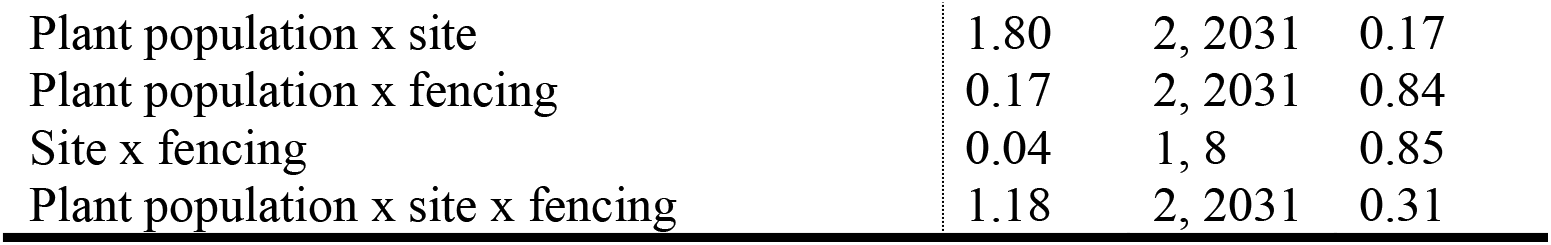
ANOVA of the effects of population, site, and fencing on (a) flowering time, (b) likelihood of producing nodules, (c) number of nodules produced per cm root, (d) specific leaf area (SLA), and (e) height. Bold type denotes significant effects.

| **(a) Date of first flower** | F | df | *P* |
| --- | --- | --- | --- |
| Plant population | 23.34 | 2, 2073 | **<0.0001** |
| Site | 40.81 | 1, 8 | **0.0002** |
| Fencing | 59.86 | 1, 8 | **0.04** |
| Plant population x site | 10.62 | 2, 2073 | **<0.0001** |
| Plant population x fencing | 1.95 | 2, 2073 | 0.14 |
| Site x fencing | 0.39 | 1, 8 | 0.55 |
| Plant population x site x fencing | 0.19 | 2, 2073 | 0.83 |
| **(b) Likelihood of nodulation** | $\chi^2$ | df | *P* |
| Plant population | 8.43 | 2 | **0.01** |
| Site | 27.63 | 1 | **<0.0001** |
| Fencing | 0.03 | 1 | 0.85 |
| Plant population x site | 3.72 | 2 | 0.16 |
| Plant population x fencing | 1.10 | 2 | 0.58 |
| Site x fencing | 0.05 | 1 | 0.82 |
| Plant population x site x fencing | 0.63 | 2 | 0.73 |
| **(c) Nodules per cm root** | F | df | *P* |
| Plant population | 2.84 | 2, 1419 | 0.058 |
| Site | 13.71 | 1, 8 | **0.006** |
| Fencing | 0.60 | 1, 8 | 0.46 |
| Plant population x site | 0.46 | 2, 1419 | 0.63 |
| Plant population x fencing | 1.97 | 2, 1419 | 0.14 |
| Site x fencing | 0.002 | 1, 8 | 0.96 |
| Plant population x site x fencing | 1.65 | 2, 1419 | 0.19 |
| **(d) SLA** | F | df | *P* |
| Plant population | 5.26 | 2, 2095 | **0.005** |
| Site | 0.07 | 1, 8 | 0.80 |
| Fencing | 0.004 | 1, 8 | 0.95 |
| Plant population x site | 2.39 | 2, 2095 | 0.09 |
| Plant population x fencing | 0.09 | 2, 2095 | 0.91 |
| Site x fencing | 0.07 | 1, 8 | 0.80 |
| Plant population x site x fencing | 0.25 | 2, 2095 | 0.78 |
| **(e) Height** | F | df | *P* |
| Plant population | 2.74 | 2, 2031 | 0.06 |
| Site | 6.66 | 1, 8 | **0.03** |
| Fencing | 0.53 | 1, 8 | 0.49 |
| Plant population x site | 1.80 | 2, 2031 | 0.17 |
| Plant population x fencing | 0.17 | 2, 2031 | 0.84 |
| Site x fencing | 0.04 | 1, 8 | 0.85 |
| Plant population x site x fencing | 1.18 | 2, 2031 | 0.31 |

**Table S2.** ANCOVA of the effects of flowering date, nodules/cm root, specific leaf area (SLA), height, and site on residual family mean relative fitness. Bold type denotes significant effects.

|  | F | df | *P* |
| --- | --- | --- | --- |
| Date of first flower | 0.05 | 1, 196 | 0.82 |
| Nodules/cm root | 0.55 | 1, 196 | 0.46 |
| SLA | 0.19 | 1, 196 | 0.66 |
| Height | 0.40 | 1, 196 | 0.53 |
| Site | 7.86 | 1, 196 | **0.006** |
| Date of first flower x site | 0.05 | 1, 196 | 0.83 |
| Nodules/cm root x site | 0.49 | 1, 196 | 0.48 |
| SLA x site | 0.02 | 1, 196 | 0.90 |
| Height x site | 1.82 | 1, 196 | 0.18 |

**Figure S1.**
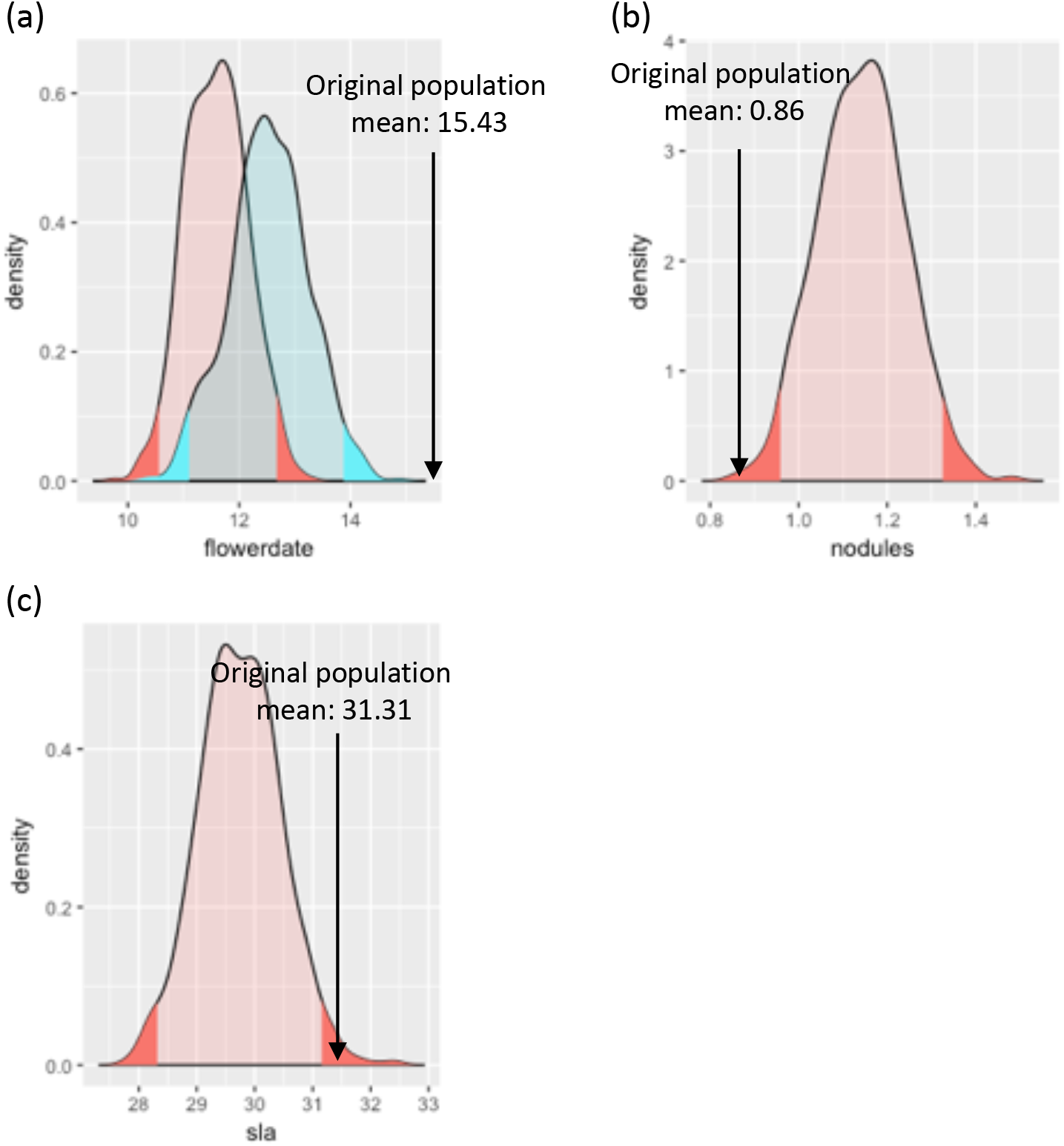
Bootstrapped trait distributions of (a) flowering time of the Lux (in red) and Marshall (in blue) populations, (b) Lux nodules/cm root, and (c) Lux specific leaf area. Original population means are shown for each trait. Shading indicates 95% confidence intervals.

**Figure S2.**
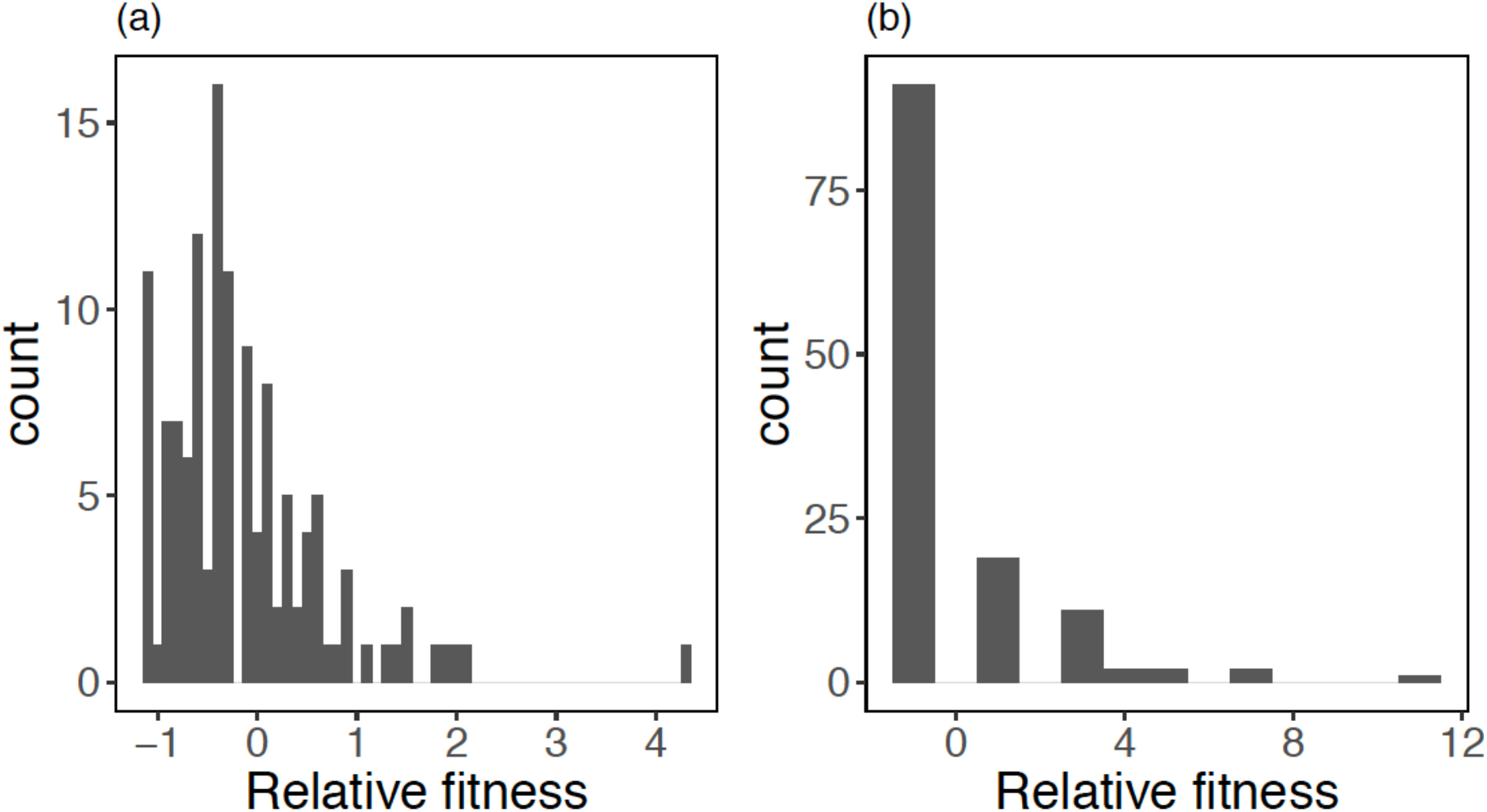
Histograms showing the distribution of residual (after accounting for spatial variation) relative fitness values of plants grown at (a) the Lux site and (b) the Marshall site.

**Figure S3.**
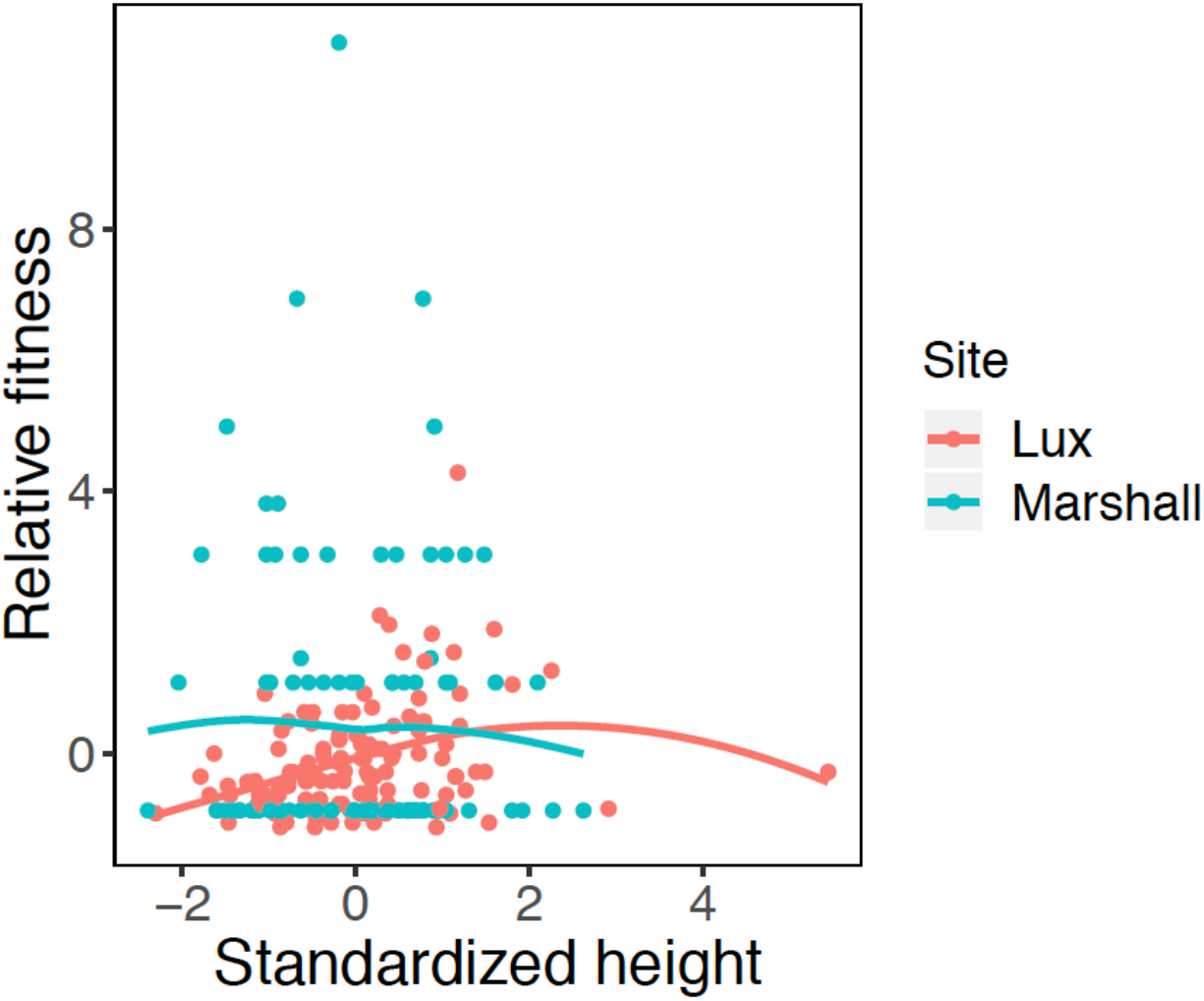
Genotypic selection on plant height at the Lux site in 2016. Graph shows the relationship between height and relative fitness, fitted with a LOESS curve.

**Figure S4.**
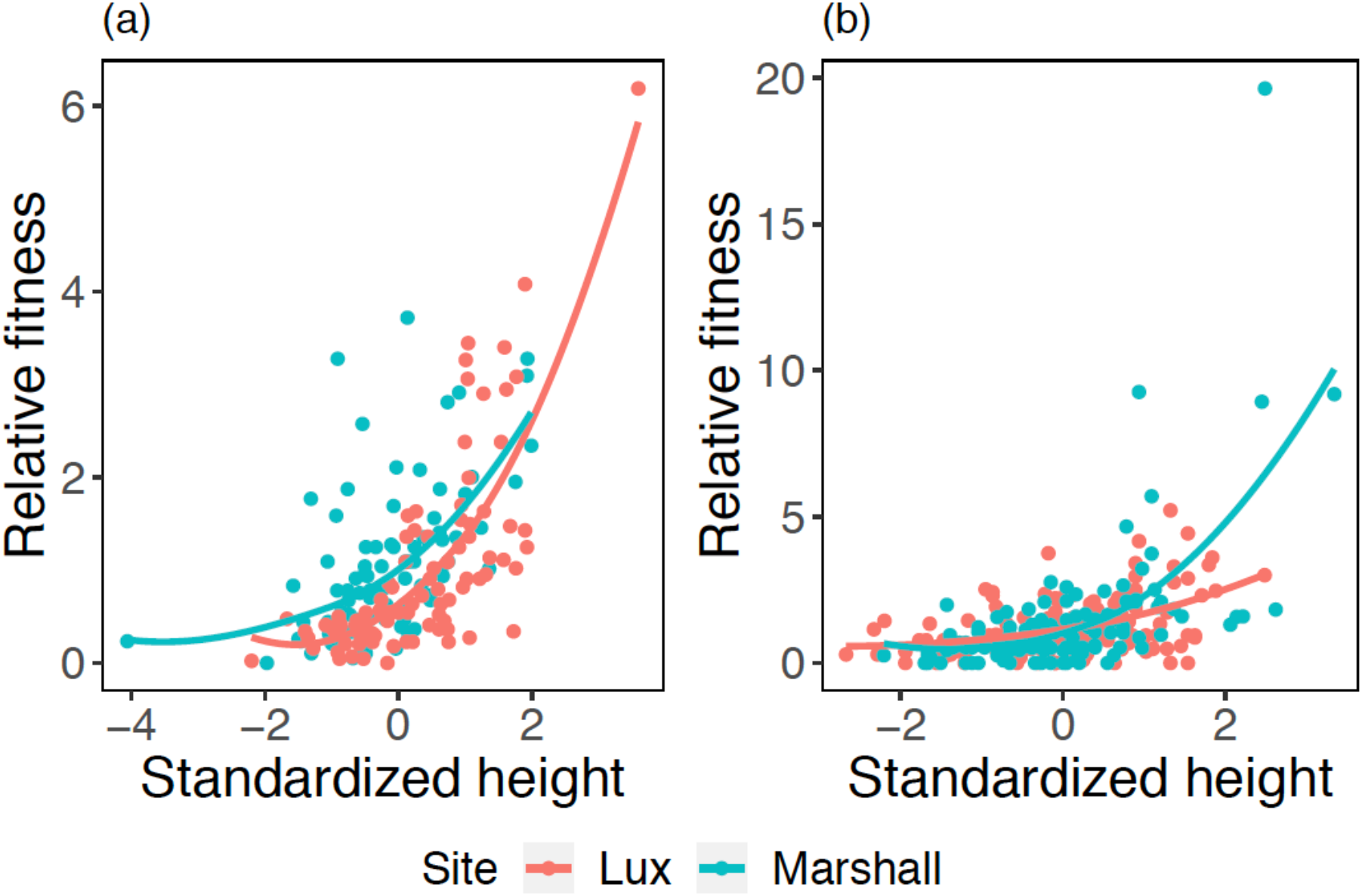
Selection on plant height at the Lux and Marshall sites in (a) 2014 and (b) 2015. Graphs show the relationship between height and relative fitness, fitted with a LOESS curve.

**Figure S5.**
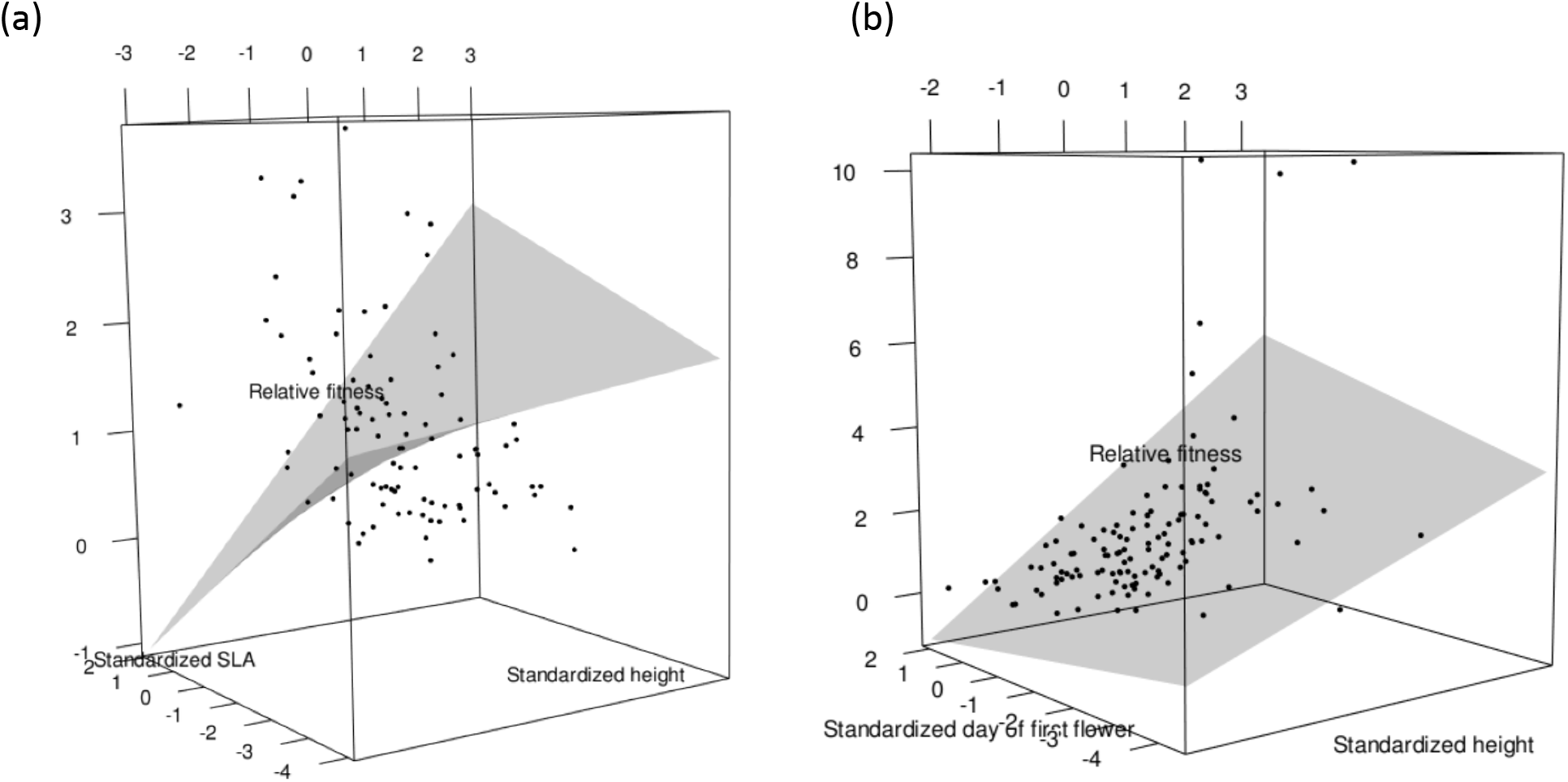
Response surfaces showing (a) the relationship between height, specific leaf area, and relative fitness and (b) the relationship between height, day of first flower, and relative fitness

